# Association between development of early cortical activity on amplitude-integrated EEG and 5-years neurodevelopmental outcome in the preterm infant

**DOI:** 10.1101/562488

**Authors:** Maria Feldmann, Valentin Rousson, Thi Dao Nguyen, Vera Bernet, Cornelia Hagmann, Beatrice Latal, Giancarlo Natalucci

## Abstract

**Aim:** To analyse the association between early aEEG and cognitive outcome at early school-age in very preterm infants.

**Methods:** Prospective cohort study including infants with gestational age (GA) <32.0 weeks, undergoing continuous aEEG recording during first 4 days of life. Semiquantitative and quantitative (maximum/minimum amplitude) measures were averaged over the recording period. Cognitive outcome was assessed with the Kaufman-Assessment Battery for Children at 5 years of age. Uni- and multivariate logistic regressions were calculated between aEEG parameters and normal cognitive outcome (IQ≥85).

**Results:** Among 118 monitored preterm children, 89 were assessed at median(IQR) corrected age of 68.6 months (65.5-71.2) [48% female, median(IQR) GA 29.9(28.2,30.9) weeks, mean(SD) birth weight 1235(363) grams]. Mean(SD) IQ was 97.8(12.7). IQ<85 occurred in 21.3 %, cerebral palsy was found in 2.2%. Despite univariate associations of total maturity scores, cycling subscores, background pattern and minimum aEEG amplitude with normal cognitive outcome none of the associations remained significant after adjustment for confounders. Socioeconomic status was identified as independent predictor of neurodevelopmental outcome.

**Conclusion:** In this cohort of very preterm infants, early short-term aEEG was not predictive of later cognitive outcome. Further research is needed to explore how aEEG could help to inform long-term prognosis in this population group.

**Key notes:**

- Preterm born infants are at high risk for neurodevelopmental impairment.
- Early amplitude integrated electroencephalography characteristics are univariately associated with cognitive outcome at 5 years of age in preterm born children.
- However, socioeconomic factors and neonatal morbidity were stronger predictors of long-term neurodevelopmental outcome than early aEEG measures.

## Introduction

Preterm born infants are at high risk for cognitive impairment in childhood and adolescence (1). During the intermediate postnatal period, very preterm infants are particularly vulnerable to brain lesions and other non-neurological complications. Amplitude integrated electroencephalography (aEEG) has been proven to be a valuable tool for cerebral function monitoring and allows for detection of background pattern deteriorations due to brain injury as well as subclinical seizures (2). The simplified set up and interpretability of aEEG enables continuous neuromonitoring and thus allows the intensive care team to immediately respond to changes in cerebral functioning (3). Beyond the guidance for clinical decision making, aEEG background pattern serves as an indicator of cerebral functional (4, 5) as well as structural (6) maturation. In other at risk populations, such as infants with hypoxic ischemic encephalopathy, aEEG background pattern is one of the most powerful predictors of neurodevelopmental outcome (7). A similar association has been found in neonates after open heart surgery (8) in which background pattern and ictal discharges on aEEG can furthermore help to detect brain lesions (9).

In the preterm population, neonatal aEEG metrics, such as cycling, background pattern, maturity score and seizure activity, have been associated with neurodevelopmental outcome up to 3 years of age (10-17). However, there is a significant heterogeneity in studies that results from a broad variety of assessed aEEG parameters as well as outcome measurements, as systematically reviewed by Fogtmann et al. (18). Additionally, whereas several studies focused on the predictive value for early childhood outcome, information on the association of early aEEG with neurodevelopmental outcome at school-age is lacking. Thus, to date, aEEG has not been broadly implemented into standard clinical neuromonitoring of preterm infants.

Hence, the aim of the present study was to assess the association of early aEEG recorded during the first 4 days of life and the trajectory of aEEG characteristics between day 1 and 4 with neurodevelopmental outcome at 5 years of age in very preterm infants.

## Methods

### Study population

This prospective cohort study was conducted in the Department of Neonatology of the University Hospital Zurich, Switzerland between January 2009 and February 2012. Inborn preterm infants with a gestational age (GA) < 32.0 weeks were included. Patients with congenital anomalies, central nervous system infections or metabolic disorders were excluded. GA was assigned based on the best obstetrical estimates according to last menstrual cycle and first trimester ultrasound if available. Cranial ultrasound was performed at days 1, 3 and 7 of life and repeated weekly until hospital discharge. Mild brain lesion were defined as intraventricular haemorrhage grade I–II and/or periventricular leukomalacia grade I, whereas severe brain lesions were defined as grade III intraventricular haemorrhage and periventricular haemorrhagic infarction, cerebellar haemorrhage, cerebral atrophy and cystic periventricular leukomalacia according to Papile et al. (19) and De Vries et al. (20), respectively. Socioeconomic status (SES) was assessed based on maternal education and paternal occupation, was scored according to Largo et al (21) and transformed so that higher scores indicate higher SES (2–12). Perinatal characteristics and type of sedation during aEEG recording were prospectively noted.

### aEEG assessment

Cerebral functional monitoring was carried out with a two-channel Brainz BRM3 aEEG monitor (Natus Medical, San Carlos, CA) and recorded with bi-parietal hydrogel electrodes, corresponding to C3 and P3 as well as C4 and P4 of the international electroencephalogram classification 10-20 system with a ground F_Z_ placed on the right or left shoulder (22). The aEEG was obtained by amplification and filtering of the raw EEG and attenuation of the activity at < 2Hz and > 15Hz was performed. The amplitudes were integrated semilogarithmicly, rectified and time compressed (1 h/6 cm of recording display scale). Continuous aEEG recording was conducted from the first to the fourth day of life. aEEG pattern analysis was performed on 3 h periods of cross-cerebral P3-P4 aEEG recordings with impedance below 12 kΩ. Tracing periods with either suspected seizure activity or artefacts were excluded from the analysis as described elsewhere (6).

### Visual and quantitative aEEG analysis

Two scoring systems were applied to semiquantitatively analyse the aEEG pattern. Brain maturation was assessed by the maturity score established by Burdjalov et al. (5). The score is composed of 4 component variables (continuity, cycling, amplitude of lower boarder and bandwidth) that sum up in a total maturity score ranging from 0 to 13 with increasing score indicating more mature brain activity. The cycling component was furthermore separately considered as “cycling subscore” with scores ranging from 0 to 5 and increasing with cycling activity (5). The background pattern was evaluated according to the classification that was introduced by Hellstrom-Westas and Rosen (23): 0 = inactive, flat (FT), 1 = low voltage (LV), 2 = burst-suppression (BSA), 3 = discontinuous (DC), 4 = continuous (C). Semiquantitative aEEG parameters were averaged for the whole recording period from day 1 to day 4. A minimum of two measurements to determine the averages were required. Quantitative analysis of the aEEG tracings was automatically performed by means of the BrainZ Analyze Research software (Chart analyser 1.71; Liggins Institute, Auckland, New Zealand). Raw aEEG data was exported and 1-min average values of the maximum and minimum amplitudes were calculated for each period. To analyse the association between quantitative aEEG measures and neurodevelopmental outcome measurements were averaged for the whole recording period. Only a single measurement was required, since aEEG min and max parameters were already averaged values. To investigate the trajectory of aEEG parameters during the first 4 days of life the slopes were calculated from a minimum of three measurements. Cohen’s kappa (95 % CI) for inter-rater agreement between the two aEEG raters (G.N., C.H.) blinded to the neonatal course was 0.79 (0.75; 0.82) for the total maturity score and 0.60 (0.52; 0.66) for the cycling subscore, respectively. For statistical analysis one rater’s aEEG scores (G.N.) were considered.

### Neurodevelopmental assessment at 5 years of age

Cognitive outcome at the age of 5 years was assessed with the German version of the Kaufman Assessment Battery for Children 1^st^ (K-ABC) (24) and 2^nd^ (K-ABC-II) edition by experienced developmental paediatricians at the Child Development Centre at the University Children’s Hospital Zurich. The mental processing composite scale of the K-ABC and the mental processing index of the K-ABC-II were considered equivalent to an intelligence quotient (IQ) with a mean (SD) value of 100 (± 15), as a general measure of cognitive ability. IQ ≥ 85 was considered as normal. Additionally, each child underwent a standardized neurological examination. Cerebral palsy was defined according Rosenbaum et al (25).

### Statistical analysis

Descriptive statistics for continuous variables were reported as median and interquartile range (IQR) as they were non-normally distributed. Merely weight at birth was normally distributed and accordingly reported as mean and standard deviation (SD). Proportions were reported for categorical variables. Unadjusted and adjusted logistic regression models were calculated to evaluate the association between either the average or the trajectory of aEEG parameters during the first four days of life with normal neurodevelopmental outcome (IQ ≥ 85) at 5 years of age. The trajectory of an aEEG parameter was summarized by the slope of a regression line calculated separately for each individual. In the adjusted model, perinatal characteristics known to influence outcome have been considered as potential confounders: GA, male sex, administration of morphine for sedation during aEEG monitoring, small for GA (i.e. birth weight < 10 centile). Furthermore, socioeconomic status (SES) has been included as covariate due to its well documented influence on neurodevelopmental outcome (21). In the logistic regression models, for the sake of comparison and without affecting any p-value, aEEG parameters were standardized such that the provided odds ratios (OR) refer to an increase of one standard deviation whatever the aEEG parameter considered. Missing values in SES were imputed by means of multiple imputation. P-values < 0.05 were considered statistically significant. All analyses were performed using R version 3.4.2 (26).

### Ethics

The institutional ethics boards of the Canton of Zurich (KEK ZH StV-35/08) approved the study protocol. Written informed consent was obtained from the parents or primary caretakers.

## Results

### Study population

Continuous aEEG recording was performed in 120 preterm infants, 2 infants had to be excluded due to a later diagnosis of leukaemia and diagnosis of a genetic syndrome, [47.5 % female, median [IQR] GA of 29.9 [28.2, 30.9] weeks and mean (SD) birth weight of 1235 (363) grams]. Perinatal characteristics of the study participants stratified by their 5-year neurodevelopmental outcome with IQ ≥ 85 are displayed in Table 1. aEEG tracings of all infants were evaluated. The aEEG recording began at a median [IQR] age of 18 [12, 21] h and was continuously performed until 87 [81, 96] h after birth. Cranial ultrasound was mildly abnormal in 20 infants (16.9 %) and severely abnormal in 7 infants (5.9 %).

**Table 1.** Perinatal characteristics of the study infants stratified by neurodevelopmental outcome.

| Perinatal variable | Neurodevelopmental Outcome |  |
| --- | --- | --- |
|  | normal<br>*n* = 70 | abnormal<br>*n* = 19 |
| Gestational age (weeks) M [IQR] | 29.7 [28.6, 30.7] | 29.0 [26.9, 29.9] |
| Birth weight, (g) m (SD) | 1267.4 (369.4) | 1013.7 (250.2) |
| Small for gestational age, n (%) | 11 (15.7) | 3 (15.8) |
| Male sex, n (%) | 34 (48.6) | 11 (57.9) |
| Preeclampsia, n (%) | 16 (22.9) | 3 (15.8) |
| Chorioamnionitis, n (%) | 15 (21.4) | 4 (21.1) |
| Antenatal steroids, n (%) | 65 (92.9) | 18 (94.7) |
| Cesarean section, n (%) | 62 (88.6) | 17 (89.5) |
| Arterial cord pH, M [IQR] | 7.3 [7.3, 7.4] | 7.3 [7.2, 7.4] |
| 5-min Apgar, M [IQR] | 7.5 [6.0, 9.0] | 6.0 [6.0, 8.0] |
| SNAPPEII, m (SD) | 19.7 (17.0) | 27.3 (19.0) |
| Respiratory distress, n (%) | 52 (75.4) | 14 (77.8) |
| Surfactant, n (%) | 18 (26.1) | 11 (61.1) |
| Days on ventilation M [IQR] | 0.0 [0.0, 1.0] | 2.0 [1.0, 6.0] |
| Major brain lesions, n (%) | 2 (2.9) | 1 (5.3) |
| Caffeine during aEEG, n (%) | 15 (21.4) | 8 (42.1) |
| Indomethacin during aEEG, n (%) | 19 (27.1) | 8 (42.1) |
| Morphine during aEEG, n (%) | 6 (8.6) | 7 (36.8) |
| SES, M [IQR] | 9.0 [8.0, 12.0] | 8.0 [5.8, 8.2] |
| CP, n (%) | 0 (0.0) | 2 (10.5) |

M = Median; m = mean; IQR = interquartile range, SD = standard deviation; Small for gestational age, i.e. birth weight < 10 centile; SNAPPE-II, Score for Neonatal Acute Physiology with Perinatal Extension-II; SES, socioeconomic status; CP, cerebral palsy.

### Neurodevelopmental outcome at 5 years of age

Neurodevelopmental follow up at 5 years has been performed in 89 infants [4 infants died, 25 were lost to follow up (Follow up rate 79 %)] at a median [IQR] corrected age of 68.6 [65.5, 71.2] months. Baseline characteristics differed between infants with and without follow up. Subjects that were lost to follow up had significantly higher GA and birth weight as well as less often respiratory distress (Supplementary Table 1). Follow up assessment at 5 years revealed a mean (SD) IQ of 97.8 (12.7) ranging from 65 to 121. IQ < 85 occurred in 21.3 %, whereas cerebral palsy was found in 2.2 %.

### aEEG measures in children with normal versus abnormal outcome

Neonates with abnormal neurodevelopmental outcome at 5 years of age had lower scores for aEEG background pattern (median [IQR] 2.8 [2.4, 3.6] versus 3.4 [3.0, 3.9]), total maturity score (3.8 [2.0, 6.4] versus 6.0 [3.5, 8.5]) and cycling subscore (0.7 [0.0, 1.4] versus 1.3 [0.5, 2.2]). Of the quantitative parameters, the minimum aEEG amplitude (mean (SD) 4.7 (1.4) versus 5.3 (1.0)), as well as the maximum aEEG amplitude were lower (12.0 (3.9) versus 13.1 (2.5)) in infants with abnormal compared to normal neurodevelopment when averaged over the 4 days of recording period.

### Prediction of outcome by aEEG measures

In univariate logistic regression analyses we found that higher total maturity scores, cycling subscores, as well as more mature background pattern and a higher minimum aEEG amplitude were associated with increased odds of normal cognitive outcome at 5 years of age. After adjusting for the confounders GA, sex, sedation, small for GA and SES none of the associations remained significant (Table 2, Figure 1). The results of the adjusted logistic regression models showed that each unit increase of SES was associated with a 40 % increase in the odds of having an IQ ≥ 85. Furthermore, ‘morphine sedation during aEEG monitoring’ was associated with a decrease in the odds of having a normal neurodevelopmental outcome with ORs ranging from 0.1–0.2. This indicates that the odds of having an IQ ≥ 85 was 5-10 times lower for children that required sedation during the first 4 days of life. When comparing perinatal clinical characteristics of sedated versus non-sedated neonates we found that sedated neonates had lower GA (p = 0.003) and birth weight (p = 0.009), higher rate of respiratory distress (p = 0.022), required more often surfactant (p < 0.0001) as well as mechanical ventilation (p < 0.0001). The Score for Neonatal Acute Physiology with Perinatal Extension-II (27) (SNAPPE-II) was also higher in sedated versus non-sedated neonates (p = 0.041) (Supplementary Table 2).

**Table 2:**
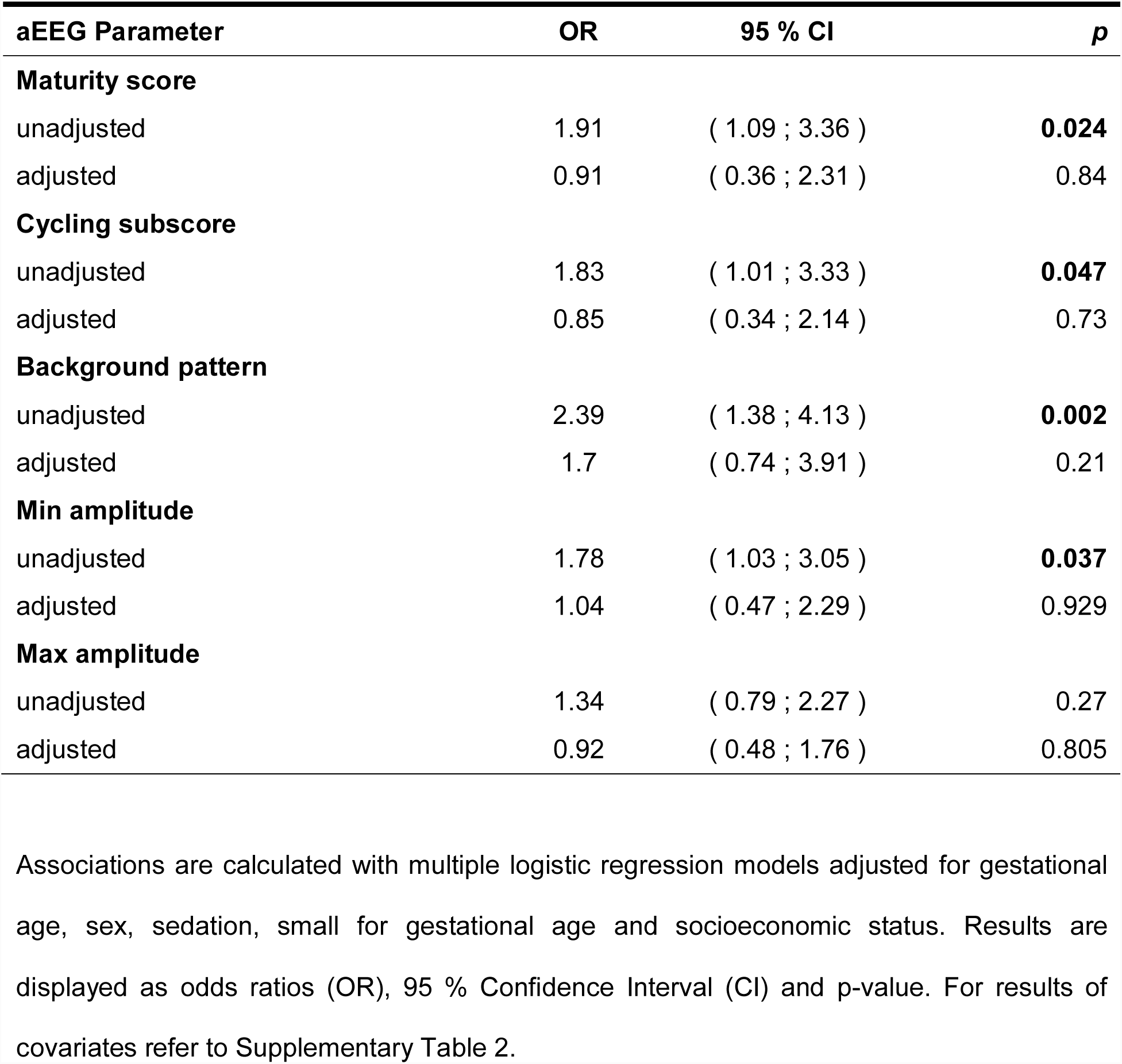
Unadjusted and adjusted association between aEEG measures and cognitive outcome at 5 years of age.

**Figure 1:**
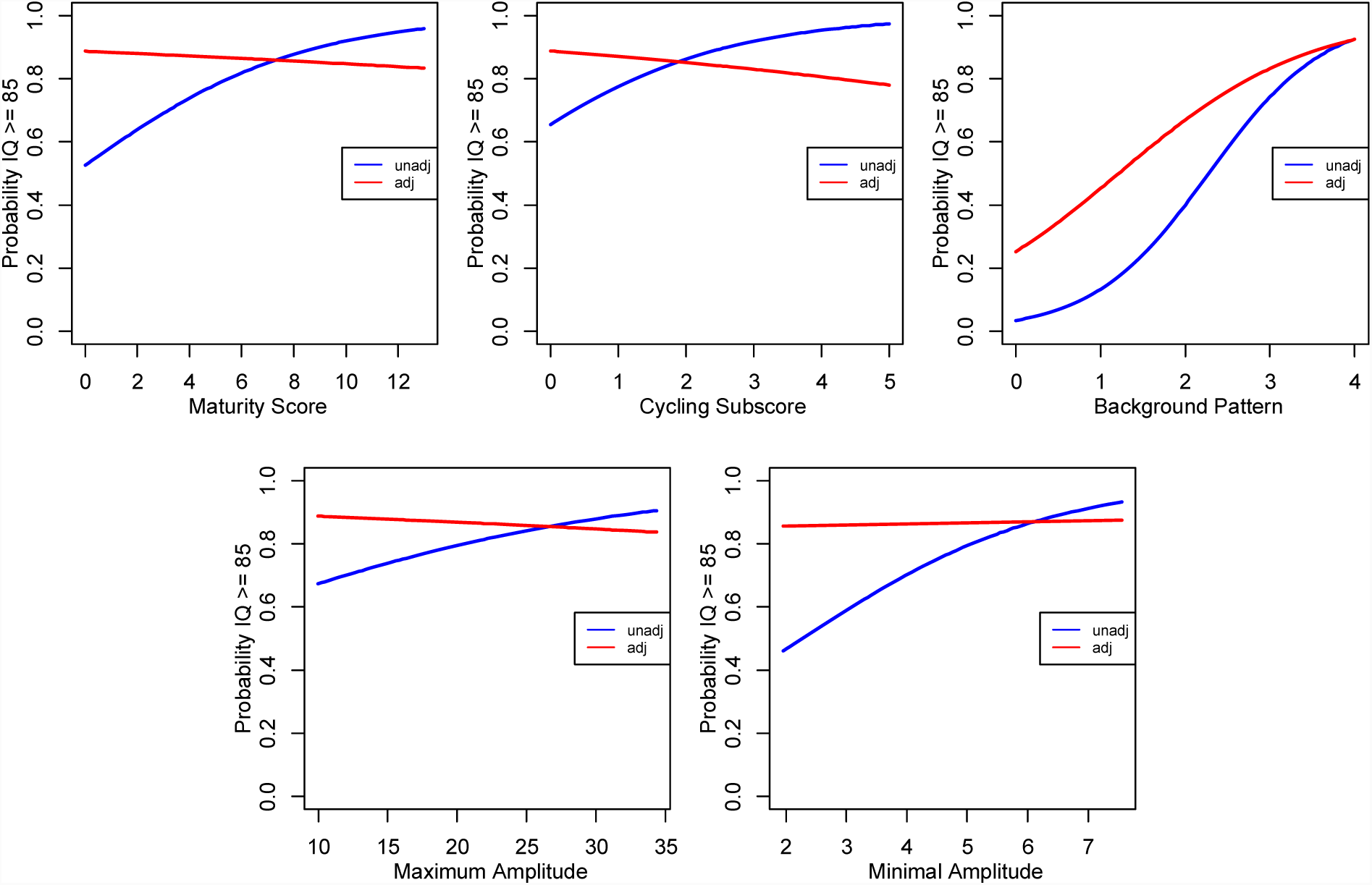
Simple and multiple linear regression models of semiquantitative (upper panel) and quantitative aEEG parameters (lower panel) and normal neurodevelopmental outcome at 5 years of age. Blue unadjusted model. Red adjusted for gestational age, socioeconomic status, sex, morphine sedation during aEEG and small for gestational age. Unadj, unadjusted; Adj, adjusted.

### Prediction of outcome by aEEG trajectories

We also examined whether the trajectories of aEEG, quantified as the slope of aEEG characteristics over the 4-day recording period, were associated with neurodevelopmental outcome. No association was found in univariate and multiple logistic regression models with neurodevelopmental outcome at 5 years of age. For details, refer to Table 3.

**Table 3:** Unadjusted and adjusted association between slopes of aEEG measures trajectories and cognitive outcome at 5 years of age.

| **aEEG Parameter Slopes** | **OR** | **95 % CI** | ***p*** |
| --- | --- | --- | --- |
| **Maturity score** |  |  |  |
| unadjusted | 1.87 | ( 0.79 ; 4.46 ) | 0.156 |
| adjusted | 1.33 | ( 0.52 ; 3.4 ) | 0.554 |
| **Cycling subscore** |  |  |  |
| unadjusted | 1.77 | ( 0.87 ; 3.6 ) | 0.118 |
| adjusted | 1.14 | ( 0.49 ; 2.65 ) | 0.764 |
| **Background pattern** |  |  |  |
| unadjusted | 0.97 | ( 0.57 ; 1.65 ) | 0.904 |
| adjusted | 0.92 | ( 0.51 ; 1.67 ) | 0.793 |
| **Min amplitude** |  |  |  |
| unadjusted | 1.19 | ( 0.65 ; 2.17 ) | 0.582 |
| adjusted | 1.15 | ( 0.54 ; 2.49 ) | 0.715 |
| **Max amplitude** |  |  |  |
| unadjusted | 0.83 | ( 0.49 ; 1.39 ) | 0.471 |
| adjusted | 0.85 | ( 0.45 ; 1.63 ) | 0.631 |

Association are calculated with multiple logistic regression models adjusted for gestational age, sex, sedation, small for gestational age and socioeconomic status. Results are displayed as odds ratios (OR), 95 % Confidence Interval (CI) and p-value.

## Discussion

This is the first prospective cohort study investigating the predictive value of early short-term quantitative and semiquantitative aEEG parameters for cognitive outcome at school age in very preterm born infants. We hypothesized that aEEG is associated with IQ at 5 years of age. Although we found evidence for univariate associations of early aEEG parameters during the first 4 days of life with cognitive outcome, these associations did not remain after adjustment for confounders. In multiple logistic regression models, we found that higher SES was independently associated with higher odds of having a normal IQ at 5 years of age. Furthermore, we found that morphine sedation was associated with a decrease in the odds of having an IQ within the normal range.

Despite previous reports of the predictive value of early aEEG recordings for neurodevelopmental outcome up to 3 years of age, we were not able to confirm the association for cognitive outcome at 5 years of age. This might be due to substantial differences in the assessment and analyses methods of aEEG recordings, analysed aEEG time points as well as outcome measurements and compositions of combined outcomes. In line with our aEEG assessment methods Bruns et al. (17) and Huning et al. (28) used the total maturity score by Burdjalov et al.(5) and the background pattern assessment by Hellstrom-Westas et al. (2), whereas others (12) applied different aEEG background scoring systems (29) as well as quantitative analysis methods such as interburst intervals (13) and dichotomized aEEG tracings into normal and abnormal (12, 14) patterns. This substantial variability in aEEG analysis methods hampers the comparability among studies and with our results. However, the aim of our study was to assess whether early aEEG recordings could inform prognosis in very preterm infants in a clinically applicable way, thus the ease of the assessment was taken into account when choosing the aEEG parameters and time points (average over 4 days) for our study. Furthermore, the absence or sparse correction for confounders applied in other studies (12, 13, 17) might explain the differences in results. In line with this we found a univariate association between the assessed aEEG parameters with neurocognitive outcome at 5 years of age, but found it to be confounded by neonatal clinical and demographic covariates such as SES and sedation.

We found that higher SES was independently associated with higher odds of having a normal IQ at 5 years of age. We chose to correct for SES, since it has been demonstrated by others to be a strong predictor of developmental trajectories in preterm born infants (21, 30). The present study results underline the importance of taking SES into consideration when analysing prediction of biomarkers for intellectual outcome in newborns at high neurodevelopmental risk, particularly on the long-term. Furthermore, in our cohort morphine sedation during aEEG monitoring was found to be associated with a decrease in the odds of having an IQ in the normal range. These results are in line with previous studies reporting that morphine administration in preterm born infants may affect aEEG tracings (31) and may be associated with adverse neurodevelopmental outcome as well as reduced regional cerebral growth (32). However, post hoc analyses of baseline characteristics among neonates who received morphine sedation during aEEG recording in our cohort compared to non-sedated neonates revealed some collinearity of ‘morphine sedation’ and other variables indicating high clinical instability of the sedated subjects (e.g. lower GA, higher SNAPPE-II score). Hence, suggesting that sedation in the present cohort is rather a surrogate marker for severity of clinical course than a predictor for adverse neurodevelopmental outcome. As we did not collect data on cumulative dose of morphine administration during the neonatal period, we are not able to analyse any further association between neonatal morphine exposition and neurodevelopment as already reported for instance by Zwicker et al. (32).

The early time-point and the brevity of the monitoring period might give another explanation for the absence of a predictive value of early aEEG with 5-year outcome in the present cohort. Relevant neonatal morbidities might occur later than during the first week of life and are not mirrored by early aEEG background pattern deteriorations. Among them, some clinical complications such as necrotizing enterocolitis, bronchopulmonary dysplasia, or sepsis have a well-described impact on neurodevelopmental outcome (33). The complexity of outcome prediction might be even more pronounced for school-age compared to early infancy outcome and could explain the differing results of our study and previous reports. In that sense the repetition of aEEG recording at term equivalent age might be more informative in terms of long term prognosis of neurodevelopmental outcome as recently demonstrated by El Ters et al. (34).

This study has some limitations that need to be mentioned. The study presents the predictive value of early aEEG in a cohort with favourable outcome. Mean IQ in our cohort was in the normal range, however IQ < 85 and cerebral palsy occurred at a higher rate than in the normal population. Although intellectual outcome in our cohort was predominantly favourable it is in line with results from previous studies as systematically reviewed by Brydges et al. (1). We investigated the association of semiquantitative and qualitative aEEG analyses methods with neurodevelopmental outcome at 5 years of age and did not assess other event-based quantitative aEEG analyses approaches, such as spontaneous activity transients frequency and length of interburst intervals, that have been shown by others to be associated with brain growth (35) and might thus be predictive of outcome. Brain injury was rare in our cohort, thus analyses of the predictive value of aEEG in subgroups according to brain injury severity was not possible.

### Conclusion

In this prospective cohort study, we found a univariate association between early semiquantitative and quantitative aEEG measures and cognitive outcome at early school-age in very preterm born children. However, socioeconomic factors and neonatal morbidity were stronger predictors of long-term neurodevelopmental outcome than early aEEG measures.

## Acknowledgments

We gratefully thank all children and their parents who participated in this study.

## Abbreviations

aEEG: amplitude-integrated electroencephalography
GA: gestational age
IQ: intelligence quotient
IQR: interquartile range
K-ABC: K-ABC-II, Kaufman Assessment Battery for Children 1st and 2nd edition
m: mean
M: median
OR: odds ratio
SD: standard deviation
SES: socioeconomic status
SNAPPE-II: Score for Neonatal Acute Physiology with Perinatal Extension-II

---

**Supplementary Table 1:**
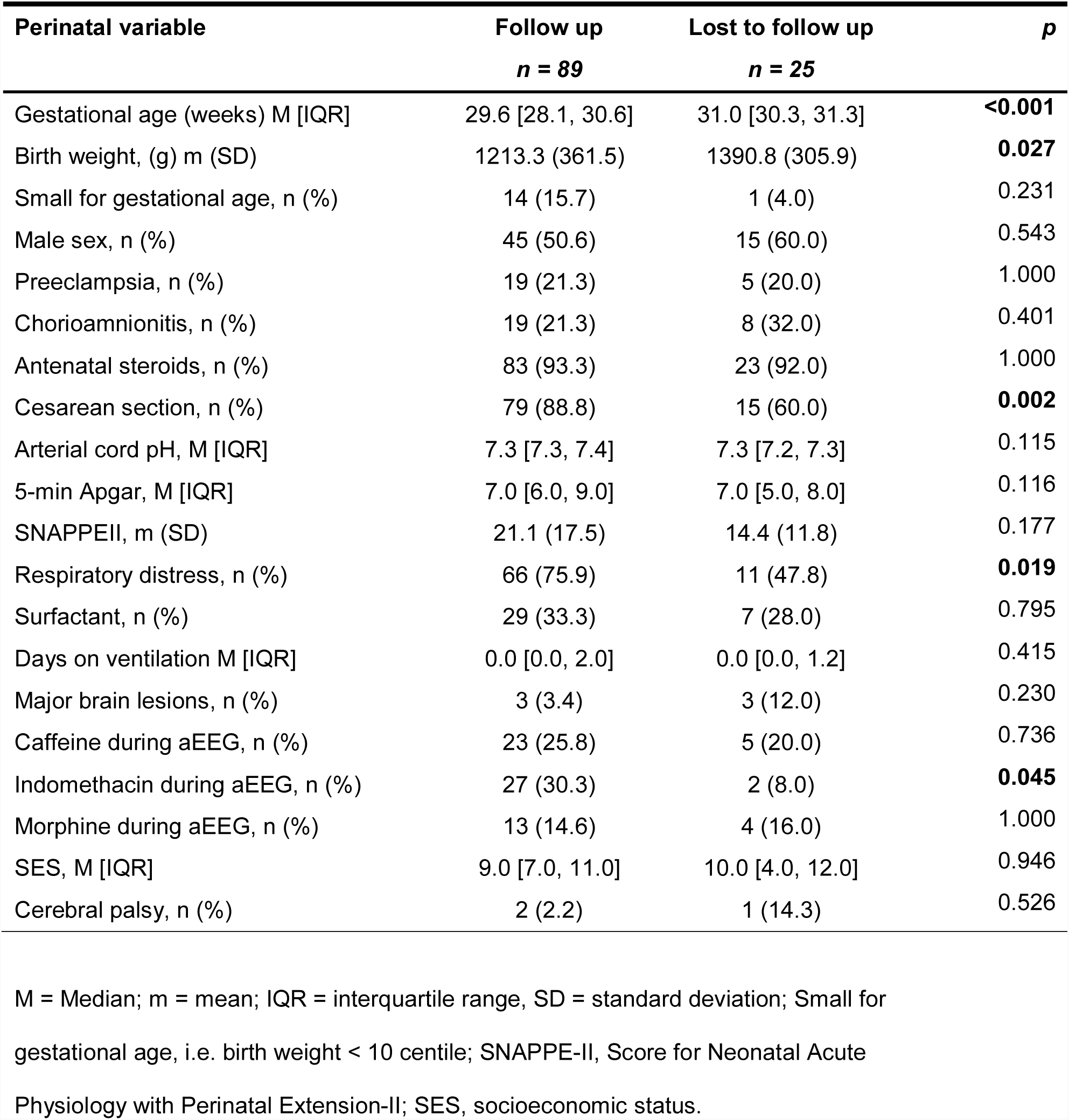
Perinatal characteristics of the study infants with follow up assessment compared to subjects that were lost to follow up.

**Supplementary Table 2:** Results of complete multiple logistic regression models for association of aEEG measures with cognitive outcome at 5 years of age.

| **Covariates** | **OR** | **95 % CI** | **p** |
| --- | --- | --- | --- |
| **Maturity score** | 0.91 | ( 0.36 ; 2.31 ) | 0.84 |
| GA | 1.54 | ( 0.95 ; 2.51 ) | 0.08 |
| Male | 0.42 | ( 0.11 ; 1.56 ) | 0.194 |
| Sedation | 0.14 | ( 0.02 ; 0.81 ) | **0.028** |
| Small for GA | 0.97 | ( 0.18 ; 5.16 ) | 0.97 |
| SES | 1.43 | ( 1.1 ; 1.87 ) | **0.008** |
| **Cycling subscore** | 0.85 | ( 0.34 ; 2.14 ) | 0.73 |
| GA | 1.58 | ( 0.96 ; 2.59 ) | 0.072 |
| Male | 0.42 | ( 0.11 ; 1.57 ) | 0.199 |
| Sedation | 0.14 | ( 0.03 ; 0.71 ) | **0.017** |
| Small for GA | 1 | ( 0.19 ; 5.41 ) | 0.998 |
| SES | 1.43 | ( 1.09 ; 1.88 ) | **0.009** |
| **Background pattern** | 1.7 | ( 0.74 ; 3.91 ) | 0.21 |
| GA | 1.3 | ( 0.84 ; 2.03 ) | 0.242 |
| Male | 0.47 | ( 0.13 ; 1.75 ) | 0.259 |
| Sedation | 0.3 | ( 0.05 ; 1.71 ) | 0.175 |
| Small for GA | 0.94 | ( 0.18 ; 4.97 ) | 0.943 |
| SES | 1.43 | ( 1.1 ; 1.86 ) | **0.007** |
| **Minimal amplitude** | 1.04 | ( 0.47 ; 2.29 ) | 0.929 |
| GA | 1.45 | ( 0.92 ; 2.29 ) | 0.11 |
| Male | 0.41 | ( 0.11 ; 1.55 ) | 0.19 |
| Sedation | 0.19 | ( 0.04 ; 0.95 ) | **0.044** |
| Small for GA | 1.32 | ( 0.2 ; 8.84 ) | 0.774 |
| SES | 1.44 | ( 1.1 ; 1.87 ) | **0.007** |
| **Maximum amplitude** | 0.92 | ( 0.48 ; 1.76 ) | 0.805 |
| GA | 1.48 | ( 1 ; 2.21 ) | 0.053 |
| Male | 0.4 | ( 0.11 ; 1.5 ) | 0.174 |
| Sedation | 0.17 | ( 0.04 ; 0.81 ) | **0.027** |
| Small for GA | 1.42 | ( 0.21 ; 9.65 ) | 0.72 |
| SES | 1.43 | ( 1.1 ; 1.87 ) | **0.008** |

Results are displayed as odds ratios (OR), 95 % Confidence Interval (CI) and p-value. GA, gestational age; SES, socioeconomic status.

